# Remodeling of the maternal gut microbiome during pregnancy is shaped by parity

**DOI:** 10.1101/2020.12.04.412205

**Authors:** Alexander SF Berry, Meghann K Pierdon, Ana M. Misic, Megan C. Sullivan, Kevin O’Brien, Ying Chen, Robert N Baldassano, Thomas D Parsons, Daniel P Beiting

**Affiliations:** Department of Pathobiology, School of Veterinary Medicine, University of Pennsylvania, Philadelphia, PA, USA; Department of Clinical Studies - New Bolton Center, School of Veterinary Medicine, University of Pennsylvania, Philadelphia, PA, USA; Department of Pediatric Gastroenterology Hepatology and Nutrition, Children’s Hospital of Philadelphia, Philadelphia, PA 19104

**Keywords:** Pig, Pregnancy, Parity, Gut Microbiome, Neonate Microbiome, Early-life microbiota, 16S rRNA sequencing, Shotgun Metagenomics

## Abstract

**Background:** The maternal microbiome has emerged as an important factor in gestational health and outcome and is associated with risk of preterm birth and offspring morbidity. Epidemiological evidence also points to successive pregnancies – referred to as maternal parity – as a risk factor for preterm birth, infant mortality, and impaired neonatal growth. Despite the fact that both the maternal microbiome and parity are linked to maternal-infant health, the impact of parity on the microbiome remains largely unexplored, in part due to the challenges of studying parity in humans.

**Results:** Using synchronized pregnancies and dense longitudinal monitoring of the microbiome in pigs, we describe a microbiome trajectory during pregnancy and determine the extent to which parity modulates this trajectory. We show that the microbiome changes reproducibly during gestation and that this remodeling occurs more rapidly as parity increases. At the time of parturition, parity was linked to the relative abundance of several bacterial species, including *Treponema bryantii, Lactobacillus amylovorus*, and *Lactobacillus reuteri*. Strain tracking carried out in 18 maternal-offspring ‘quadrads’ – each consisting of one mother sow and three piglets – linked maternal parity to altered levels of *Akkermansia muciniphila, Prevotella stercorea*, and *Campylobacter coli* in the infant gut 10 days after birth.

**Conclusions:** Collectively, these results identify parity as an important environmental factor that modulates the gut microbiome during pregnancy and highlight the utility of a swine model for investigating the microbiome in maternal-infant health. In addition, our data show that the impact of parity extends beyond the mother and is associated with alterations in the community of bacteria that colonize the offspring gut early in life. The bacterial species we identified as parity-associated in the mother and offspring have been shown to influence host metabolism in other systems, raising the possibility that such changes may influence host nutrient acquisition or utilization. These findings, taken together with our observation that even subtle differences in parity are associated with microbiome changes, underscore the importance of considering parity in the design and analysis of human microbiome studies during pregnancy and in infants.

## Introduction

The mammalian microbiome plays a key role in maternal and infant health, and recent studies have highlighted the value of the maternal microbiome for predicting the risk of preterm birth [1–3], the leading cause of neonatal death worldwide [4]. Although the exact mechanisms by which maternal microbes might influence pregnancy and offspring health have yet to be fully defined, studies in mice have begun to provide clues. For example, microbial metabolites produced in the maternal gut can be detected in the placenta and fetal tissues, where they drive postnatal innate immune development [5]. Similarly, short chain fatty acids produced by the maternal microbiome cross the placenta, where they signal through multiple host pathways to protect offspring from metabolic disease [6]. Microbes are also vertically transmitted to offspring during birth and in the perinatal period, and these early colonizers can have long-term effects on child development [7–9]. Despite the growing recognition that the maternal microbiome influences infant health, we currently have a remarkably poor understanding of the clinical and environmental factors that can impact the microbiome during pregnancy.

Maternal parity – the number of previous pregnancies – is associated with increased risk of preterm birth in humans [10, 11], but studies of parity and health in humans are often confounded by numerous socioeconomic and psychosocial factors [12]. Since both the maternal gut microbiome and parity have been identified as key determinants of gestational health, it is important to understand whether parity influences the microbiome during pregnancy. Previous studies in dairy cows have shown that animals pregnant for the first time (nulliparous) have different uterine and rumen microbiome compositions than do animals with only a single prior pregnancy (primiparous) or two or more previous pregnancies (multiparous)[13, 14]. However, it is as yet unclear if parity impacts either the maternal gut microbiome during pregnancy or the microbiome of the developing offspring.

Elucidating the relationship between microbiome composition and pregnancy in human subject research is challenging. Studies are often characterized by small sample sizes [15], cross-sectional or sparse longitudinal sampling [16], and present challenges in controlling for confounding factors. Interpersonal variation in diet has a particularly large impact on gut microbiota composition in human studies [17], and differences in maternal diet during pregnancy have been shown to influence the infant gut microbiome [18]. Properly controlling for diet often involves either retrospective studies paired with self-reported food intake surveys [19], highly controlled feeding studies [20], or a focus on geographically distinct populations that differ in dietary practices [21–25], all of which are made more challenging in pregnancy. In addition, studies of pregnancy and the microbiome in humans rarely address parity due to limited numbers of pregnancies in most countries. Large animal models offer an appealing alternative to human studies for examining parity. Pigs are commonly used as biomedical models of humans due to similarities in anatomy and physiology [22–24], and have provided valuable insight into functions of the human gut microbiome [22–24]. Here, we describe high-resolution microbiome profiling during synchronized pregnancy in sows, carried out in a highly controlled environment, to determine if the microbiome changes during pregnancy, and whether parity plays a role in this process.

16S rRNA marker gene sequencing and shotgun metagenomics were used to assess the association between pregnancy and the gut microbiome in a population of mother sows with parity ranging from zero to seven, where diet and environment are meticulously controlled. Maternal fecal samples collected weekly throughout the 114-day gestation, together with samples from piglets born to these mothers, allowed direct comparisons to be made between maternal and infant microbiomes. We observed that 1) the maternal gut microbiome changes predictably during pregnancy, 2) this remodeling occurs more rapidly in high parity animals, compared to their low parity counterparts, 3) parity is associated with gut microbiome composition at parturition, and 4) the composition of the early infant gut microbiome is influenced by parity. Taken together, our results highlight the importance of considering parity in maternal gut microbiome studies and suggest that pregnancy history can shape both the maternal microbiome as well as early colonization events in offspring, albeit likely by different mechanisms.

## Results

### Predictable changes in microbiome composition throughout pregnancy in the pig

Previous studies examining the microbiome in pigs have focused either on broad stages of gestation [26], or on post-natal growth and feed efficiency [27–30]. In order to understand the extent to which the maternal gut microbiome is affected during pregnancy in pigs, stool samples were collected weekly from mother sows beginning at gestational day 34, when pregnancy was first confirmed by ultrasound, and throughout the full 114-day gestation. The resulting 390 stools samples (10-12 per sow) were subjected to microbiome profiling by targeted sequencing of the V4 region of the 16S rRNA gene. In order to leverage the longitudinal aspect of this data, we trained a supervised regression model on all samples from 60% of the animals to calculate a microbiome maturity index during pregnancy [31, 32], then tested the model on the remaining animals. This approach is ideal for longitudinal data because it quantifies the relative rate of change in microbiome composition over time. Our model identified a significant correlation (*P*<3.3e-13; *R*^2^=0.27) between the actual versus predicted day of gestation **(Fig 1A)**. Despite a trend toward overestimating day of gestation early in pregnancy and underestimating later, our model accurately predicted which samples belonged to late-versus early-term pregnancies. The families *Porphyromonadaceae* and *Muribaculaceae* were the most important taxa for predicting day of gestation **(Fig 1B)**. To identify transitions in microbial community structure during pregnancy, we applied a Dirichlet multinomial mixtures (DMM), which uses a probabilistic approach to carry out unsupervised clustering of taxonomic data into community types. This analysis showed that the gut microbiome of most sows occupies Cluster 1 from the time that pregnancy is confirmed until day 65 of gestation, indicating a shared microbiota structure early in pregnancy **(Fig 1C)**. However, as animals moved through pregnancy, the gut microbiome underwent a shift marked by a departure from Cluster 1, evident by day 72. **(Fig 1C)**. *Treponema and Clostridium sensu stricto* were the most important taxa for clustering gut microbiota communities by DMM **(Fig 1D, Fig 1E)**.

**Figure 1.**
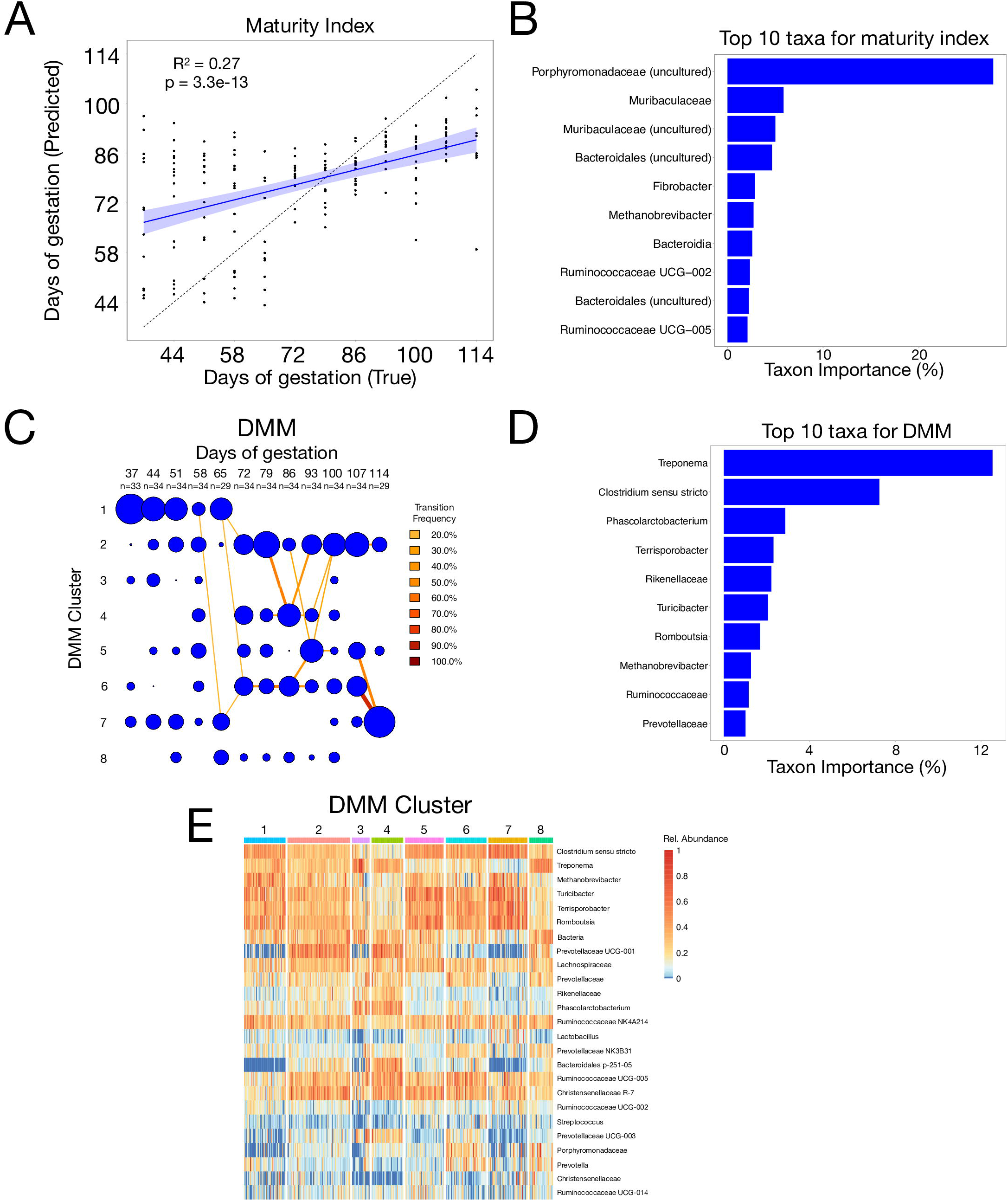
Gut microbiota compositional changes during gestation follow several predictable trends. 16S sequencing of fecal samples from 34 mother sows sampled longitudinally during the 114-day gestation reveals several trends in gut microbiota changes. A) A maturity index trained on 60% of the animals and tested on the remaining 40% shows that the amount of time of gestation can be predicted (*P*=3.3e-13) with some accuracy (*R*^2^=0.27) using gut microbiota composition data. B) The 10 taxa that contributed the most to the accuracy of the maturity index are shown in order of importance. C) Dirichlet Multinomial Mixtures (DMM) bins samples into one of 8 clusters, each defined by a unique gut microbiota composition. Days of gestation is on the X-axis, and clusters are labeled on the Y-axis. The total number of samples at each time point is also labeled on the X-axis. The size of the blue circle is proportional to the number of samples contained in each cluster. Transitions made by a substantial percentage of individuals (Transition Frequency >20%) are indicated by lines connecting blue clusters. D) The 10 taxa that contributed the most to the accuracy of the DMM are shown in order of importance. E) Heatmap shows the relative abundance of taxa (normalized across each taxa) for each sample. Samples are grouped by DMM cluster to allow for visualization of the taxa that are enriched or reduced on average in each DMM cluster.

### Parity modulates the pregnancy-induced remodeling of the gut microbiome

Although our maturity index and DMM analyses show a clear predictable change in the microbiome during pregnancy, there was still unexplained variation, as evident from the maturity index *R*^2^=0.27. We hypothesized that parity may contribute to some of this variation. To test this hypothesis, we grouped animals based on whether they had no prior pregnancies (zero parity, or nulliparous) (*n* animals = 9, *n* samples = 107), 1-3 prior pregnancies (low parity) (*n* animal = 13, *n* samples = 150), or 4-7 prior pregnancies (high parity) (n animals = 12, n samples = 133). A non-parametric microbial interdependence test (NMIT) was used to summarize data across all timepoints into a single microbiome trajectory value of each individual sow. Principal Coordinate Analysis (PCoA) **(Fig 2A; SFig 1)** and NMDS **(Fig 2B)** of these data showed that NMIT distances were significantly different between parity groups **(Fig 2C)**. The greatest difference in microbiome trajectory occured between zero parity (nulliparous) and low parity sows, suggesting that having even one prior pregnancy was sufficient to impact microbiome trajectory during future pregnancies. The difference in microbiome trajectories between low and high parity sows was also significant (*Adj. P*<0.05), albeit less substantial than the difference between nulliparous and low parity animals.

**Figure 2.**
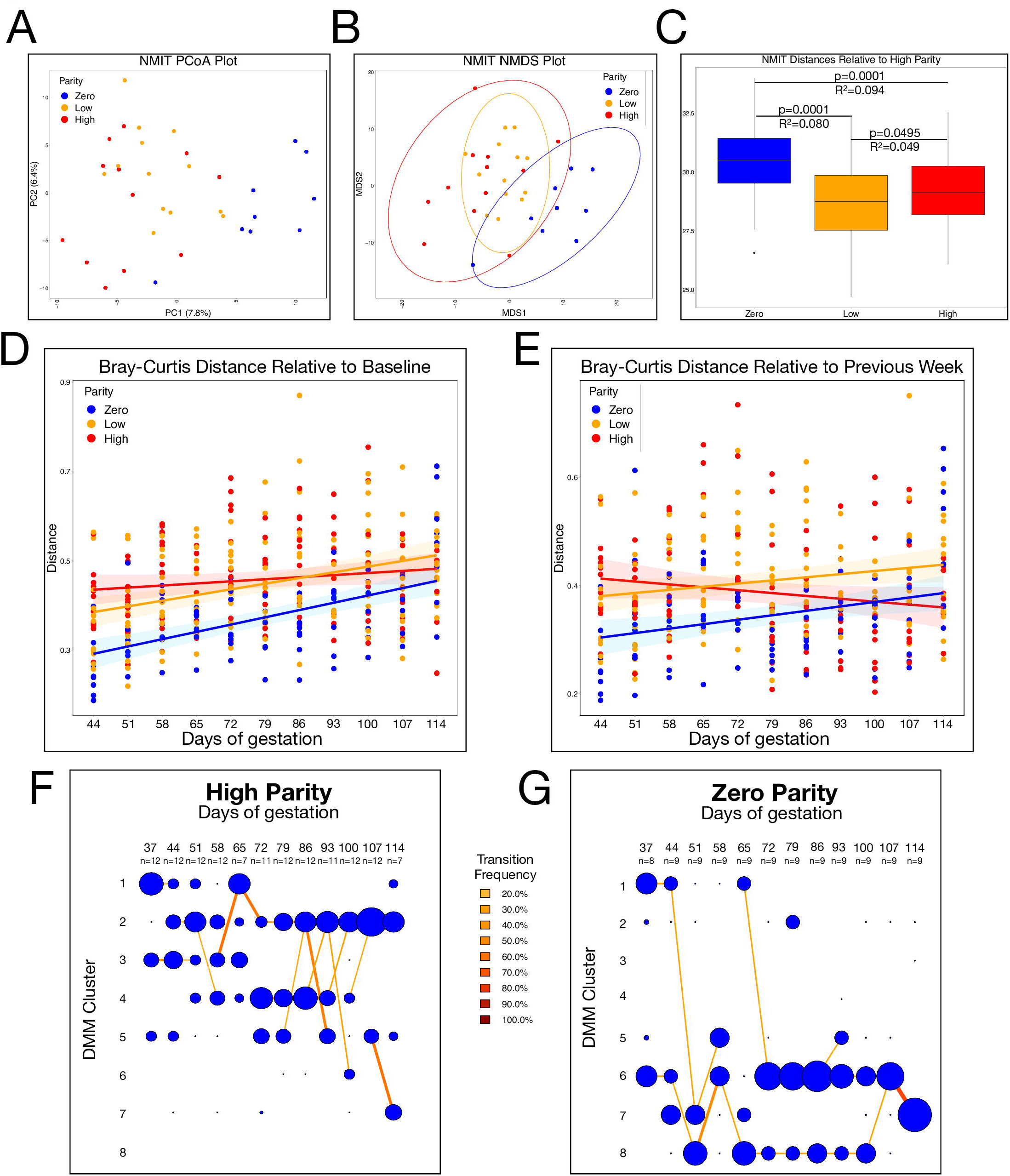
Parity affects the gut microbiota trajectory during gestation. Clustering each of the 34 individuals by parity (Zero = no previous pregnancies, Low = 1-3 previous pregnancies, High = 4-7 previous pregnancies) reveals that parity explains a significant amount of the variation between gut microbiota trajectories during gestation. A) Non-parametric microbial interdependence test (NMIT), which calculates correlations between each pair of taxa for each individual over time, was performed for each individual. A Principal Coordinate Analysis (PCoA) plot of the NMIT data shows how the gut microbiota trajectory differs between animals of different parities across the first two axes. B) Non-metric multidimensional scaling (NMDS) plot of the NMIT data depicts the differences between the gut microbiota trajectories, with 95% confidence intervals displayed as ellipses around each parity bin. C) Boxplots depict the differences in NMIT between each individual and each high parity individual. Boxplots show the median and the first and third quartiles, with whiskers that extend to outliers up to 1.5 times the interquartile range. The adjusted *P*-values and *R*^2^are shown for each comparison. D) Bray-Curtis beta diversity was calculated between each sample and the Day 37 sample from the same individual. E) Bray-Curtis beta diversity was calculated between each sample and the previous week’s sample from the same individual. F) Dirichlet Multinomial Mixtures (DMM) showing only samples collected from high parity animals. G) DMM showing only samples collected from zero parity animals.

To determine how the microbiome trajectory differs between parity groups, Bray-Curtis beta diversity between each sample and the baseline (day 37) sample from the same animal was calculated. Both Bray-Curtis **(Fig 2D)** and weighted UniFrac **(SFig 2A)** showed that nulliparous animals, compared to their low and high parity counterparts, exhibit a more subtle but incremental change from baseline as pregnancy progresses. In contrast, high parity animals underwent a more abrupt change from baseline early in pregnancy with little or no further change evident throughout. Since comparing each time point with a single baseline reference likely fails to capture the magnitude of changes that occur between time points, we could not rule out that high parity sows might undergo substantial changes from week to week throughout pregnancy while remaining consistently different from baseline. To reconcile these interpretations, beta diversity was calculated between each sample and the previous sample from the same individual using Bray-Curtis **(Fig 2E)** and weighted UniFrac **(SFig 2B)**. This analysis confirmed that the rate of change of gut microbiota composition in zero and low parity animals increases steadily throughout pregnancy, while high parity animals actually show a decrease over time, suggesting that as animals have more pregnancies, remodeling of the gut microbiome occurs more rapidly.

Since the largest differences in microbiome trajectory occur between zero and high parity animals, we further investigated the differences between these two groups over time by repeating the DMM modeling, but with animals grouped by parity. While most animals occupy the cluster 1 community type at day 37, regardless of whether they are high **(Fig 2F)** or zero parity **(Fig 2G)**, high parity sows quickly move out of cluster 1 to occupy clusters 2, 3, 4 and 5 throughout most of the pregnancy, with most of these animals occupying cluster 2 at full-term. Interestingly, nulliparous sows also leave cluster 1 early in pregnancy but occupy clusters 6, 7 and 8 instead (most in cluster 7 at full-term) – three clusters rarely, if ever, occupied by high parity sows at any point during pregnancy **(Fig 2F and 2G)**. Taken together, the data show that parity influences the gut community types that develop during pregnancy.

### Parity is associated with an altered microbial environment during the perinatal period

Our DMM analysis suggested that by the end of gestation, high parity animals had a community type that was distinct from that of zero parity animals. To better understand how parity influenced the microbial environment as animals approached parturition, we carried out shotgun metagenomic sequencing on a subset of the animals profiled above, along with their offspring. In total, 18 mother-offspring ‘quadrads’ – each comprising a mother sow and three of her piglets – were examined, including stool from 7 nulliparous sows and 11 high parity sows at baseline (day 37) and at the end of gestation (day 114), along with rectal swabs collected from 54 piglets at day 10 of life. Bray-Curtis beta diversity was calculated among high parity sows and between high and zero parity sows at both timepoints. At day 37, parity is only slightly correlated with beta diversity (*P*=0.050; *R*^2^ =0.11**)(Fig 3A)**. Conversely, parity is strongly correlated with beta diversity by day 114 (*P*=9e-4; R^2^ =0.21) **(Fig 3B)**. To identify the species driving these community composition differences, differentially-abundant taxa were calculated using linear discriminant analysis (LEfSe) at both timepoints. Consistent with the beta diversity analysis, only two low-abundance taxa were differentially-abundant between zero and high parity animals at day 37: *Selenomonas bovis* and *Prevotella copri*, both of which were more abundant among zero parity animals compared to those of high parity **(Fig 3C)**. At day 114, six taxa were differentially-abundant between zero and high parity including *Methanobrevibacter* (unclassified), Porcine type C oncovirus, *Peptostreptococcaceae* (unclassified), Treponema *bryantii, Lactobacillus amylovorus*, and *Lactobacillus reuteri* **(Fig 3D)**. The differences in beta diversity and differentially-abundant taxa at each timepoint suggest that the impact of parity on microbiome composition is modest early in pregnancy, and that parity-associated taxa emerge later as animals near parturition.

**Figure 3.**
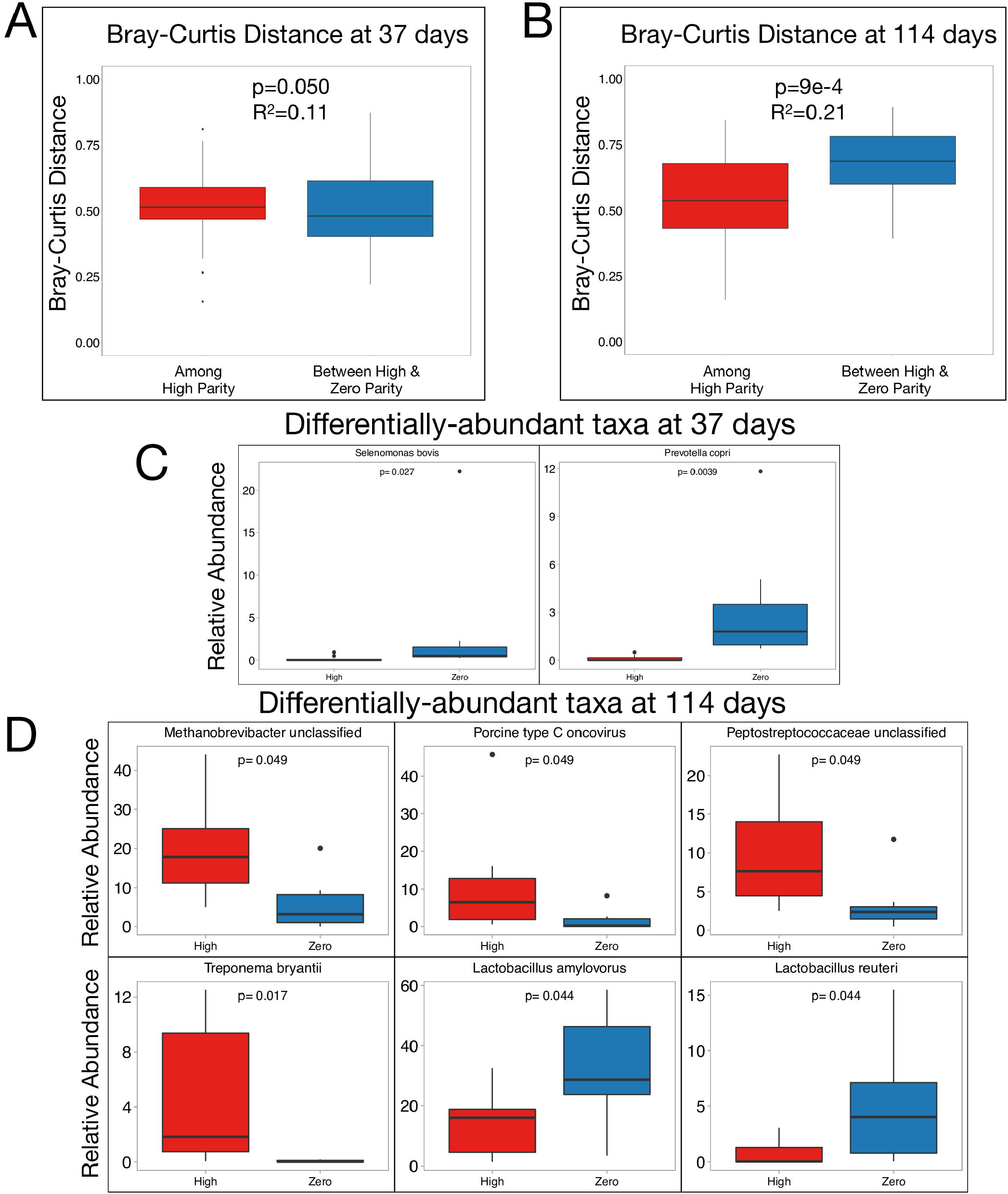
Parity is associated with significant differences in the relative abundance of key taxa at the end of gestation. Fecal samples from 18 mother sows (7 zero parity and 11 3-7 parity) were collected at Days 37 and 114 of gestation and shotgun metagenomic sequencing was performed. Bray-Curtis beta diversity was calculated between each sample and each high parity sample at **A)** the beginning of gestation (Day 37 of gestation), and **B)** just prior to delivery (Day 114 of gestation). All microbes with average relative abundance >1% across all 36 samples (Adj. *P*<0.05) that were differentially-abundant between zero and high parity animals at **C)** Day 37 and **D)** Day 114 are shown. Boxplots show the median and the first and third quartiles, with whiskers that extend to outliers up to 1.5 times the interquartile range. *P*-values (adjusted for multiple testing in C and D) and *R*^2^ are shown where appropriate.

### Maternal gut microbiome composition does not predict early infant gut colonization

Our observation that the sow gut microbiome was influenced by parity, together with recent evidence suggesting that maternal gut microbes are likely candidates for vertical transmission to offspring [33, 34], prompted us to characterize the relationship between maternal and offspring gut microbes. As expected, Bray-Curtis beta diversity analysis of metagenomic data from piglets born to each sow (n=54) showed that maternal gut microbiomes were markedly distinct from those of the piglet gut (*P*=1e-5, *R*^2^=0.27), such that the gut microbiome composition of a sow is more similar to that of other sows than to that of her infant offspring **(SFig 3A)**. Surprisingly, there was no relationship between the relative abundance of the dominant species observed in the maternal sow gut and those in their piglets. Many microbes present in the piglet gut were completely absent from the maternal gut and vice versa **(SFig 3B)**, and a linear regression comparing the relative abundance of the eight most abundant bacterial taxa in the maternal gut at Day 114 to the same taxa in the offspring gut showed no correlation (Adj. *P*>0.3) **(SFig 4)**. To explore this in more detail we focused on *Escherichia coli*, which was present at high relative abundance in both sows and piglet, and performed strain tracking using StrainPhlAn [35] to determine whether mother and offspring harbored the same strain. After removing samples for which multiple *E. coli* strains were detected and samples for which no *E. coli* was detected, a neighbor-joining phylogeny confirmed that *E. coli* strains observed in siblings are more closely related than E. coli observed in non-siblings, but also revealed that a mother and her offspring harbor distinct strains of *E. coli* **(SFig 5)**. Taken together, these data suggest that the maternal gut is unlikely to be the main source of early colonizers of the piglet gut.

**Figure 4.**
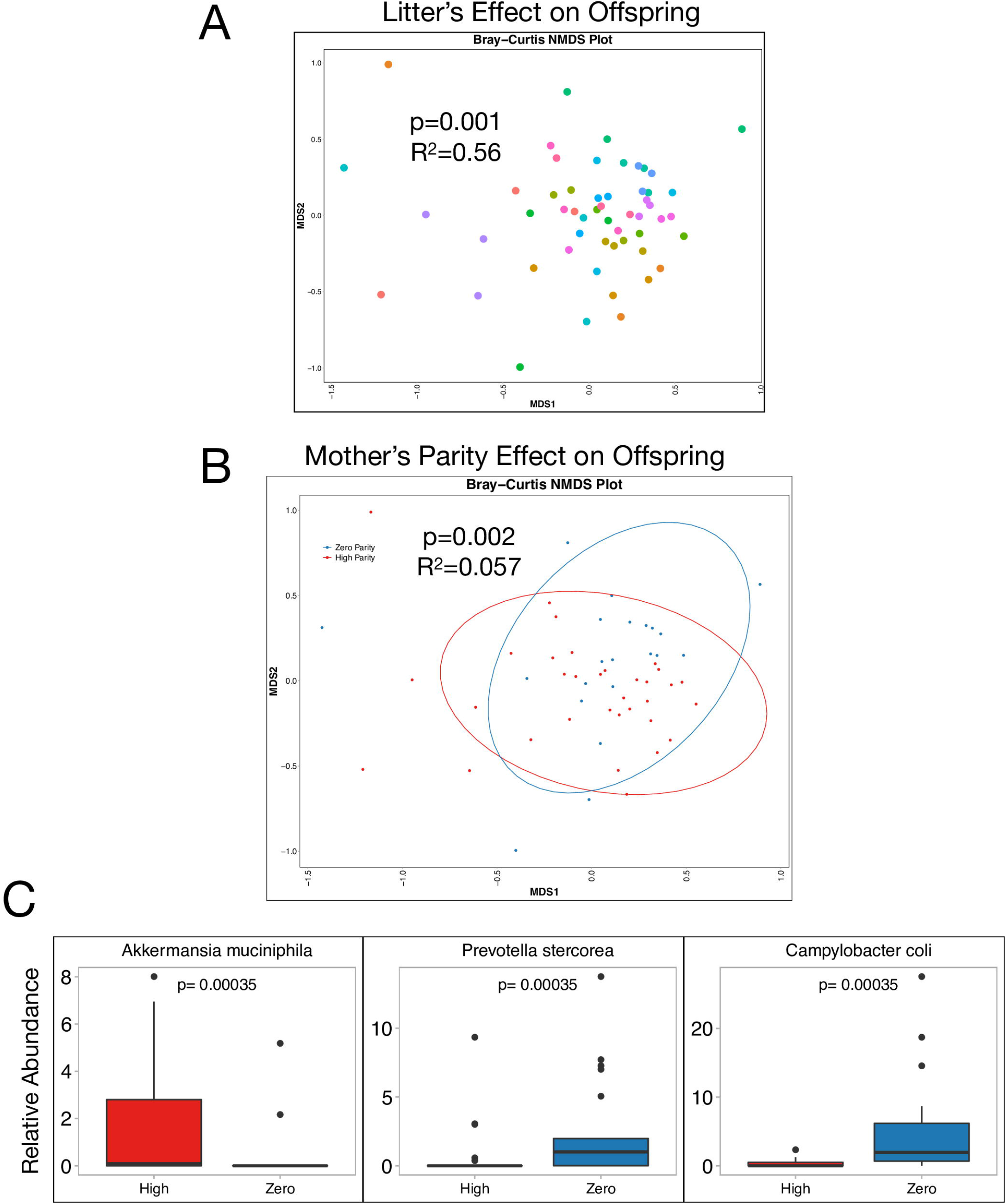
Maternal parity is associated with significant differences in offspring gut microbiome composition. Fecal swabs from 3 offspring of each of the 18 mother sows were collected 10 days after delivery and shotgun metagenomic sequencing was performed. **A)** An NMDS plot shows the differences in gut microbiome composition across all 54 piglets, as determined by Bray-Curtis distance. Piglets born to the same mother (representing a unique “litter”) are represented by the same color. Litter is significantly associated with gut microbiome composition (*P*=0.001) and explains most of the variation in gut microbiome composition (*R*^2^=0.56). **B)** The NMDS plot comparing the Bray-Curtis dissimilarity between piglet gut microbiome samples is colored by the parity of the piglet’s mother. Maternal parity is significantly associated with gut microbiome composition (*P*=0.002) and explains 5.7% of the variation in gut microbiome composition (*R*^2^=0.057). **C)** All microbes with average relative abundance >1% across all 54 piglet fecal samples (Adj. *P*<0.05) that were differentially-abundant between those born to zero and high parity mothers are shown. Boxplots show the median and the first and third quartiles, with whiskers that extend to outliers up to 1.5 times the interquartile range. Adjusted P-values are shown for each differentially-abundant taxa.

### Maternal parity is associated with altered microbiome composition in offspring

Our analysis of *E. coli* strains in metagenomic data from piglets suggested similarities in gut microbiomes of animals from the same litter. Indeed, Bray-Curtis beta diversity comparing all 54 piglets showed that litter – or mother to which a piglet is born – is significantly associated with gut microbiome composition 10 days after birth and explains more than half of the variation between piglet gut microbiomes (*P*=0.001, *R*^2^ =0.56) **(Fig 4A)**. These data indicate that, although few taxa are shared between mother and offspring in our study, maternal factors are still a major driver of the offspring microbiome. We therefore hypothesized that maternal parity could influence the offspring microbiome in ways that are independent of direct vertical transmission of taxa. Consistent with this notion, maternal parity was also significantly associated with piglet gut microbiome composition (*P*=0.002, *R*^2^ =0.057) **(Fig 4B)**. The differences between offspring born to nulliparous versus high parity mothers was driven by three bacterial taxa: *Akkermansia muciniphila, Prevotella stercorea*, and *Campylobacter coli* **(Fig 4C)**. Notably, none of these species were found to be differentially-abundant between nulliparous and high parity sows, arguing that parity impacts both maternal and offspring gut microbial communities, but that the mechanisms by which this occurs likely differs.

## Discussion

Longitudinal analysis of the gut microbiome in 34 mother sows revealed a shift in the community types present during gestation. A limited number of longitudinal studies have been conducted during pregnancy in humans, with one study reporting a dramatic shift in community composition from first to third trimester [16], while another showed remarkably stable community structure in the gut, vagina, oral cavity throughout gestation [2]. These conflicting results highlight that additional data from well-controlled animal studies, as well as from diverse human populations – such as the ongoing Multi-Omic Microbiome Study: Pregnancy Initiative (MOMS-PI) [3] – are needed to generate a more complete picture of host-microbiome interactions during pregnancy. Recent efforts to consider statistical methods for analyzing longitudinal microbiome data and carrying out normalization for cross-study comparisons [36, 37], together database efforts that enable integration of large volumes microbiome data [38], all constitute important developments that will enhance our ability to integrate and mine microbiome data from different types of maternal-infant microbiome studies.

Our data show that parity modulates gut microbiome maturation during gestation in pigs. One confounding factor in the study of parity in both our experiments as well as in human studies is age, since humans and animals of higher parity are also usually older. In addition, pigs are highly social and establish a social hierarchy based on age. Consequently, nulliparous pigs are not only the youngest animals but are also at the bottom of the social rank and would normally be subject to bullying by older animals of higher parity. To mitigate unwanted effects of social hierarchy and ensure maximum animal welfare, nulliparous and multiparous sows were separated by a fence during gestation (see methods). However, several lines of evidence argue against age being a major contributor to the differences observed in our parity analysis. First, we showed that gut microbiome composition at Day 37 of pregnancy is similar between nulliparous (younger) and multiparous (older) animals, despite the fact that these groups have been separated for approximately five weeks (Fig 3A,C). Furthermore, DMM analysis also showed that most animals, regardless of parity and age, occupy the same cluster at Day 37. Second, approximately 5 months of age typically separate sows that differ by a single parity, yet the NMIT analysis revealed that the largest change in microbiome trajectory occurs between nulliparous and single parity sows (SFig 2), rather than between the very youngest and oldest animals. Lastly, the impact of parity on the microbiome is evident even when the nulliparous (youngest) animals are excluded – high and low parity animals are not separated by the fence, yet they still differ in microbiome maturation during pregnancy (Fig 2, SFig 1B).

Despite the fact that anaerobic bacteria identified in our piglet gut samples, including *Bacteroides* species and the sulfate reducer *Disulfovibrio piger* [39], are common members of the gut microbiome, our paired metagenomic analysis of maternal-offspring ‘quadrads’ suggested that the piglet neonatal gut microbiome does not originate from the maternal gut (SFig 3-5). However, we examined piglets only at a single time point, and the reference-based metagenomic analysis we used may lack the sensitivity to detect low abundance organisms, thus we cannot not rule out the possibility that some early colonizers of the infant gut were inherited from the maternal gut. Nevertheless, the apparent lack of shared strains or species between the maternal gut microbiome at parturition and the piglet gut at day 10 of life suggests that early gut colonizers are acquired, at least in part, from other maternal or environmental sources. Interestingly, numerous *Lactobacillus* species as well as *Clostridium clostridioforme* were detected in the piglet gut, but were absent from the sow gut, and have been observed in human vaginal microbiome studies [40, 41]. *Subdoligranulum*, a genus found in nearly all our piglets, has been observed in breast milk microbiomes [42]. In humans, breastfeeding is a source of early infant gut microbes [43], and is a significant predictor of early childhood gut microbiota [44] and immunity [45]. Parity has been shown to affect the lipid and protein content in human breast milk [46], as well as the microbiota composition of cow colostrum [47]. Taken together, these data suggest that the vaginal mucosa and breast milk should be explored for their potential as early sources of microbial colonization of the piglet gut, and our piglet data raise the possibility that one or both sites may be influenced by parity.

We identified several parity-associated bacterial species in the maternal gut at the end of gestation, as well as in the piglet gut. Among these were *Treponema*, the top taxon in our DMM model of the gut microbiome during gestational (Fig 1D), and specifically the species *T. bryantii* which was identified as enriched in high parity pigs by metagenomic sequencing (Fig 3D). Although little is known about this spirochete, *T. bryantii* was recently linked to increased feed efficiency in sows [48]. *Lactobacillus* levels peaked in DMM Cluster 7 (Fig 1E), the cluster occupied by most nulliparous animals at the end of gestation (Fig 2G). Two species, *L. reuteri and L. amylovorus*, were enriched among nulliparous animals (Fig 3D), and both are common probiotic strains associated with weight and fat gain, and enhanced immune function in sows and piglets [49–52]. We also found that piglets born to high parity sows had increased relative abundance of *Akkermansia muciniphila* (Fig 4C), a species recently identified in human breast milk and breast tissue [53] and which has been widely linked to reduced risk of obesity and metabolic disease in humans [54, 55] and more recently to reduced adiposity in pigs [56]. Collectively, these data support a model whereby parity-associated changes in the microbiome have the potential to alter maternal or infant metabolism.

Given the causal role for the microbiome in a wide range of human diseases, our data suggest that the microbiome merits further exploration as a possible contributing factor to parity-associated outcomes in maternal and infant health. For example, it is well-established that parity reduces the risk for estrogen receptor- and progesterone receptor-positive breast cancers [57–59] and ovarian cancer [60], but increases the risk of dental disease [12] and dementia [61]. Although the mechanisms underlying these parity effects remain unclear, studies point to long-term alterations in hormones and systemic inflammatory mediators possible contributors [62–64]. Just as we observed an impact of parity on the offspring gut microbiome, so too are there well-documented effects of parity on infant morbidity and mortality. Offspring born to nulliparous mothers have reduced birth weight and higher mortality rates [58], and are at increased risk of childhood obesity and metabolic disease [65]. Whether and how these parity-associated phenotypes and diseases are linked to the microbiome remains an important and unresolved question. Future studies exploring this topic will be valuable not only in improving agriculture, but also for advancing microbiome-based diagnostics and therapies to improve maternal-infant health in humans.

## Conclusions

Pregnancy history affects both the maternal gut microbiome during gestation and the infant gut microbiome postpartum. Sources other than the maternal gut are potential contributors to the early piglet microbiome. Parity influences the relative abundance of key bacterial species associated with obesity and altered metabolism, including *Akkermansia muciniphila, Treponema bryantii*, and several *Lactobacillus species*.

## Methods

### Animal husbandry and sample collection

Fecal samples for our Pig Pregnancy and Parity (P3) Microbiome study were acquired from animals housed at the Penn Vet Swine Teaching and Research Center. The facility was environmentally controlled and maintained high standards of hygiene. Study animals were selected from one of two adjacent pens of gestating sows in the barn’s gestation area. The first pen housed a group of 130 gestating parous sows (having previously birthed a litter) that were maintained in a single large dynamic group and fed by two electronic sow feeding (ESF) stations (Compident VII, Schauer Agrotronics, Prambachkirchen, Austria). The second pen housed a similar, but smaller group, of 65 nulliparous sows (not having previously birthed a litter) in a single dynamic group and were fed by a single ESF station. The smaller, younger animals were separated from the older sows to help mitigate the stress and other unwanted impact of conspecific aggression that arises in the establishment of a social hierarchy as older sows tend to bully the younger animals. The pens adjoined and created a near identical physical environment for the sows but were separated by a fence line that prevented direct nose to nose contact between groups. Both groups were subjected to similar management and husbandry practices although the larger group had access to straw bedding and an outdoor concrete loafing area. These pens provide animals with a space allowance of at least 2.0 m^2^/head. Weaned, mated animals were added to the dynamic groups weekly at 8 days post-weaning. A similar number of near-term pregnant animals (>day 110 of gestation) were removed from the gestation pens and transferred to the farrowing and lactation area of the barn. Sows gave birth in individual pens (4.1 m^2^) equipped with a hinged farrowing crate. This crate initially provided protection to the newborn piglets but was opened at 10 to 14 days post-partum to provide the sow with additional mobility. Sows in the birthing pens only came into contact with their own piglets, and the newborn piglets only contacted one another and their mother for the duration of lactation. Piglets were weaned ranging from 28 to 35 days of age. Weaned sows were moved to the breeding area of the barn and housed in individual stalls while they were bred via post-cervical artificial insemination prior to returning to the gestation pens. The birthing pens were pressure washed with hot water and detergent to remove all organic matter and then disinfected prior to refilling with another near-term sow.

All sows were fed a similar standard corn–soy diet meeting or exceeding NRC [66] standards for gestating and lactating sows with a metabolizable energy of 3,197.2 kcal/kg. The quantity of feed each sow received was based on both her stage of gestation or lactation and the animal’s body condition. Animals with less condition (skinnier) received larger quantities of feed than animals with more condition (fatter). Body condition of animals was scored at placement into the gestation pen and then reevaluated at ∼30-day intervals. Standard production metrics were collected for each sow and their litter including sow age and parity (number of previous pregnancies resulting in a birth). Sows were confirmed pregnant at ∼37 days post-mating via real-time ultrasonography. In order to determine how the gut microbiome changed throughout pregnancy, a fecal sample was collected from each pregnant sow at the time of pregnancy confirmation (37 days of gestation) and every 7 days until delivery (∼114 days of gestation), for a total of 12 fecal samples per sow. To determine the effect of maternal parity and gut microbiome composition on the gut microbiome composition of offspring, fecal swabs were taken from piglets 10 days after birth. Fecal samples and swabs were stored at -80C until DNA extraction.

### 16S rRNA gene sequencing and processing

16S rRNA gene sequencing was carried out on stool samples collected weekly from 34 sows throughout gestation. This group included 9 zero parity animals (no previous pregnancies), 13 low parity animals (1-3 previous pregnancies), and 12 high parity animals (4-7 previous pregnancies). Samples were obtained from at least 10 unique time points from each sow, with most sows represented by all 12 sampling times (weekly from Day 37 to 114), for a total of 390 total fecal samples. DNA was extracted from fecal samples using Qiagen PowerSoil DNA extraction kit. 16S rRNA sequencing was performed as described previously [67]. Briefly, the V4 region of the 16S rRNA gene was amplified using PCR using Accuprime Pfx Supermix and custom primers for 30 cycles [67]. Quantification and clean-up of post-PCR products was carried out using PicoGreen reagent and AMPureXP beads, respectively. Pooled PCR libraries were quantified and sized using a Qubit 2.0 and Tapestation 4200, respectively. 250bp paired-end sequencing was performed using an Illumina MiSeq. The QIIME2 pipeline [68] was used to process and analyze 16S sequencing data using qiime2 version 2019.7.0. Samples were demultiplexed using q2-demux and denoised using Dada2 [69]. Sequences were aligned using maaft [70] and phylogenetic trees were reconstructed using fasttree [71]. Weighted UniFrac [72] and Bray-Curtis [73] beta diversity metrics were estimated using q2-core-metrics-diversity after samples were rarefied to 10000 reads per sample, and p-values were adjusted for multiple hypothesis testing using Benjamini-Hochberg (B-H) false discovery rate (FDR) corrections [74]. Taxonomy was assigned to sequences using q2-feature-classifier classify-sklearn [75] against the Silva rRNA reference database [76, 77]. Taxa were collapsed to the genus level, when possible. OTUs with less than 0.1% average relative abundance across samples, and those present in less than half of samples, were removed.

### Longitudinal data analysis

The qiime2-longitudinal plugin was used to analyze weekly sampled microbiome data from each sow [78]. A maturity index was calculated using 60% of the samples as training data to compare the expected and predicted days of gestation using qiime longitudinal maturity-index [31, 32]. To determine whether parity affects microbiome remodeling during pregnancy, the change in Bray-Curtis and weighted UniFrac beta diversity metrics over time were determined using qiime longitudinal first-distances and qiime longitudinal linear-mixed-effects. Non-parametric microbial interdependence tests (NMIT) were performed using qiime longitudinal nmit [79]. NMIT calculates correlations between each pair of taxa for each individual over time, allowing direct, quantitative comparisons of microbial interdependence between individuals as opposed to between timepoints within an individual. Dirichlet multinomial mixtures (DMM) was used to model the relationships between microbial communities as determined by 16S sequencing using R code modified from Stewart et al., 2018 [80, 81], and DMM clusters (community types) were determined based on lowest LaPlace approximation. The proportion of samples occupying a DMM cluster at each timepoint was plotted to visualize changes in community types over time during pregnancy.

### Shotgun metagenomic sequencing and analysis

DNA extracted from fecal samples obtained from 18 of the study animals early (day 37) and late in gestation (day 114) were used for shotgun metagenomics. This subset included 7 zero parity sows and 11 high parity sows (parity ≥3) (n=36 samples). DNA was also extracted from fecal swabs collected from 3 piglets from each of the 18 sows (n=54). All DNA extractions were carried out using the Qiagen PowerSoil DNA extraction kit, and sequencing libraries (n=90) were prepared following Illumina’s Nextera XT protocol. Sequencing was performed on a NextSeq500 to generate 150bp single-end reads. Reads were trimmed using trimmomatic version 0.33 [82], and quality was confirmed using FastQC [83]. MetaPhlAn version 2.6.1 was used to determine the relative abundance of microbial taxa in each sample, collapsed to species [84]. The correlation between variables such as parity and microbiota composition was determined using PERMANOVA as implemented in the vegan package [85] in R [86]. Differentially-abundant taxa were determined using LDA Effect Size (LEfSe) [87] and p-values were adjusted for multiple hypothesis testing using B-H FDR corrections in R. Nonmetric Multidimensional Scaling (NMDS) plots comparing sow and piglet microbiome composition were generated using the vegan package in R. Plots were visualized using ggplot2 [88], patchwork [89], and ggthemes [90]. Heatmap was generated using hclust2 (available at https://github.com/SegataLab/hclust2). To determine whether species found in the piglet gut originated from the maternal gut, markers from Escherichia coli, a microbe found in high abundance across sow and piglet guts, were extracted using StrainPhlAn [35]. Samples for which multiple strains were suspected were removed from the analysis. A neighbor-joining phylogenetic tree was reconstructed with 1,000 bootstrap replicates using MEGA7 [91], and visualized using FigTree version 1.4.4 (available at http://tree.bio.ed.ac.uk/software/figtree/).

## Declarations

### Ethics approval and consent to participate

Research was completed under permits issued by the Institutional Care and Use Committee of the University of Pennsylvania as IACUC Protocol #806059 under The Guide for the Care and Use of Laboratory Animals.

### Consent for publication

All authors of this work concur with this submission, and the data presented have not been previously reported nor are they under consideration for publication elsewhere.

### Availability of data and material

All 16S and shotgun metagenomic sequencing data are publicly-available on the Sequence Read Archive (SRA) under the study accession numbers PRJNA644935 and PRJNA645191, respectively. All code and software used to process and analyze the data is available as a fully reproducible computing environment on Code Ocean (https://codeocean.com/capsule/2719244/tree).

### Authors’ contributions

MKP, TDP and DPB designed the study. MKP collected samples. MKP and TDP oversaw operations in the swine facility. ASFB, KO and YC processed the samples. AMM and MCS prepared libraries and carried out sequencing. ASFB and DPB analyzed data and wrote the paper. MKP, TDP and RNB assisted with writing and data interpretation.

## Acknowledgements

Acknowledgements

We thank Dr. Shuai Wang and Ms. Elise Krespan for providing feedback on data presentation and analyses.

## Figures and Tables

**Supplemental Figure 1. The most significant difference between gut microbiota trajectories lie between nulliparous and multiparous animals**. A Principal Coordinate Analysis (PCoA) plot of the NMIT data shows how the gut microbiota trajectory differs between animals of different parities across the first two axes. Each point represents an individual’s trajectory during gestation. Each point is a number which represents the parity, and the color of each number represents the parity bin (zero, low, or high).

**Supplemental Figure 2. Weighted UniFrac beta diversity shows that parity affects the gut microbiota trajectory during gestation**. A) Weighted UniFrac beta diversity was calculated between each sample and the Day 37 sample from the same individual. B) Weighted UniFrac beta diversity was calculated between each sample and the previous week’s sample from the same individual.

**Supplemental Figure 3. Sows and piglets have significantly different gut microbiome compositions**. Shotgun metagenomic sequencing was performed for fecal samples from 18 mother sows that were collected at days 37 and 114 of gestation, and from fecal swabs from 3 offspring of each pig collected 10 days after delivery. **A)** An NMDS plot representing the Bray-Curtis beta diversity between the 90 samples (36 sow and 54 piglet) was generated. The microbiome compositions of sows are significantly different from that of piglets (*P*=1e-5). **B)** A heatmap was generated using the 32 bacterial species with relative abundance >1% among either sows or piglets. For each sample, the heatmap shows the relative abundance of each species. Consistent with the NMDS plot, the heatmap separates into two major phylogenetic branches, with sows on the left and piglets on the right. Four sow samples cluster within the piglet branch and are noted below the heatmap.

**Supplemental Figure 4. The relative abundance of bacterial taxa in a sow does not correlate with the relative abundance of that taxa in her offspring**. Linear regression analyses was performed for each of the 8 bacterial species with average relative abundance >1% across all samples and that were present in at least two-thirds of all samples. For each piglet, the relative abundance of each bacterial species in its mother at Day 114 was plotted (X-axis) against the relative abundance of that species in the piglet 10 days after delivery (Y-axis). Fecal swabs from three piglets from each sow were sequenced. There were no significant correlations between bacterial species in mother and offspring in any of the eight species (Adj. P > 0.3).

**Supplemental Figure 5. There is no evidence that *Escherichia coli* in piglets were inherited from the maternal gut**. A neighbor-joining phylogeny was reconstructed using 34 *E. coli* strains extracted from the shotgun metagenomic sequencing data from piglets and sows likely harboring only a single *E. coli* strain. Three sets of *E. coli* from different piglet-sow sets are highlighted in orange, cyan, and red. For example, highlighted in orange, the 3 piglets born to Sow 214 share similar *E. coli* genotypes, but immediately prior to birth (Day 114), the sow’s gut harbored a very different *E. coli* genotype. Similarly, in cyan, Sow 4074 and her piglet harbor different *E. coli* genotypes. Finally, in red, three piglets of Sow 1962 harbor E. coli genotypes with a most recent common ancestor at the root of the phylogeny. E. coli was chosen because it is the microbe found in the highest abundance across both sows and piglets, thus providing the deepest coverage of strains.

