## Supplementary figures and images for "Remodeling of the maternal gut microbiome during pregnancy is shaped by parity"

### Supplemental figure 1

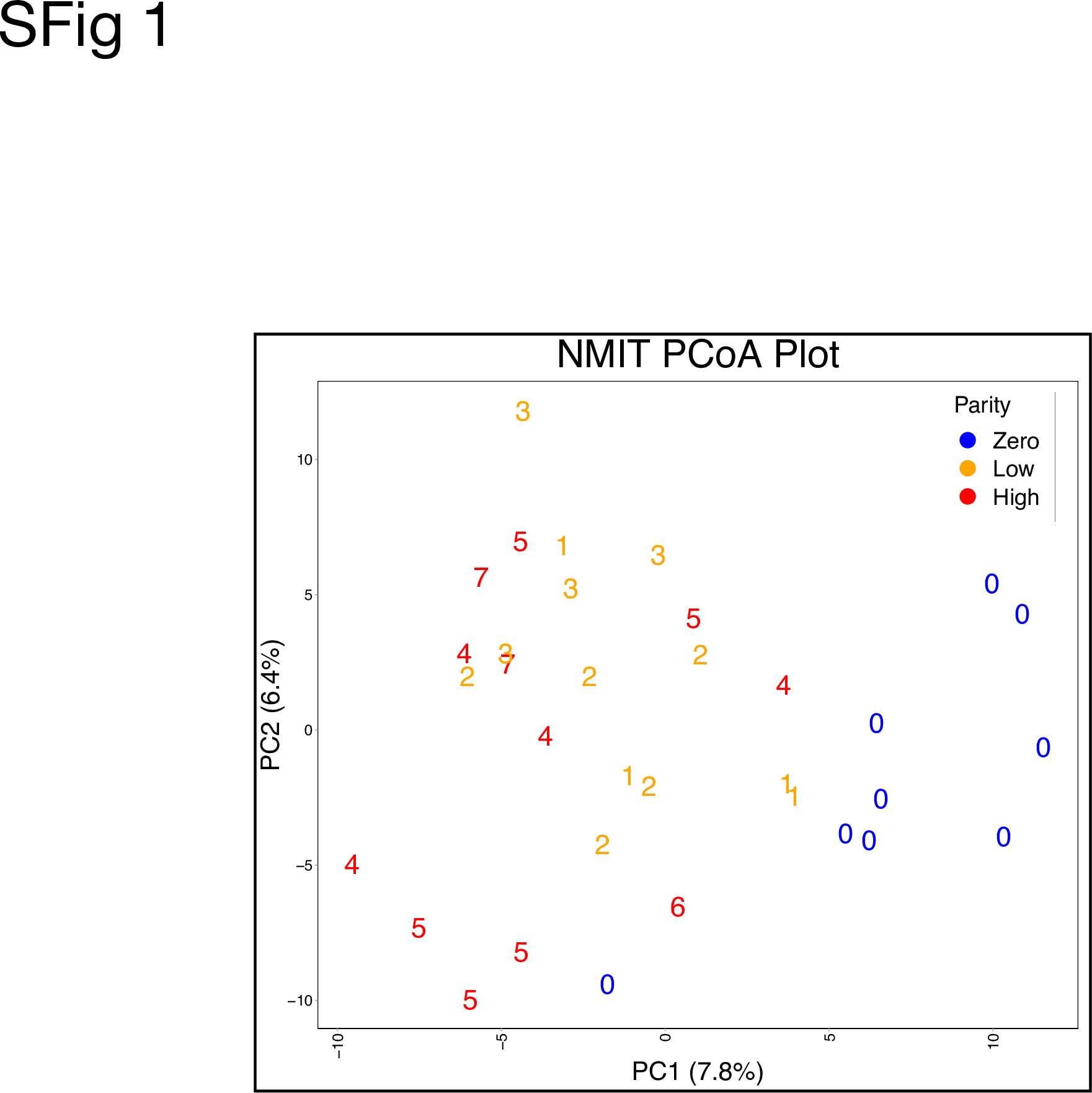

### Supplemental figure 2

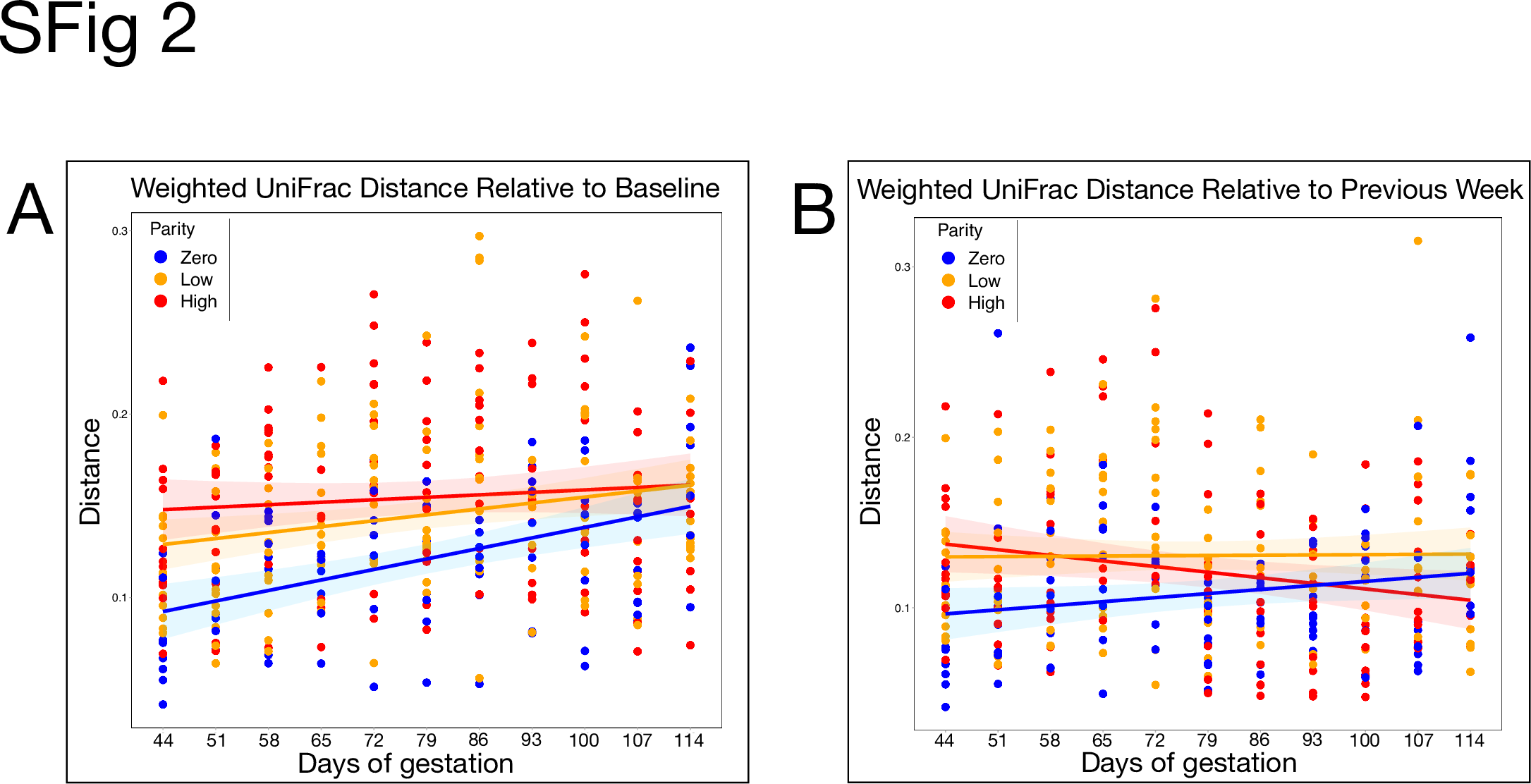

### Supplemental figure 3

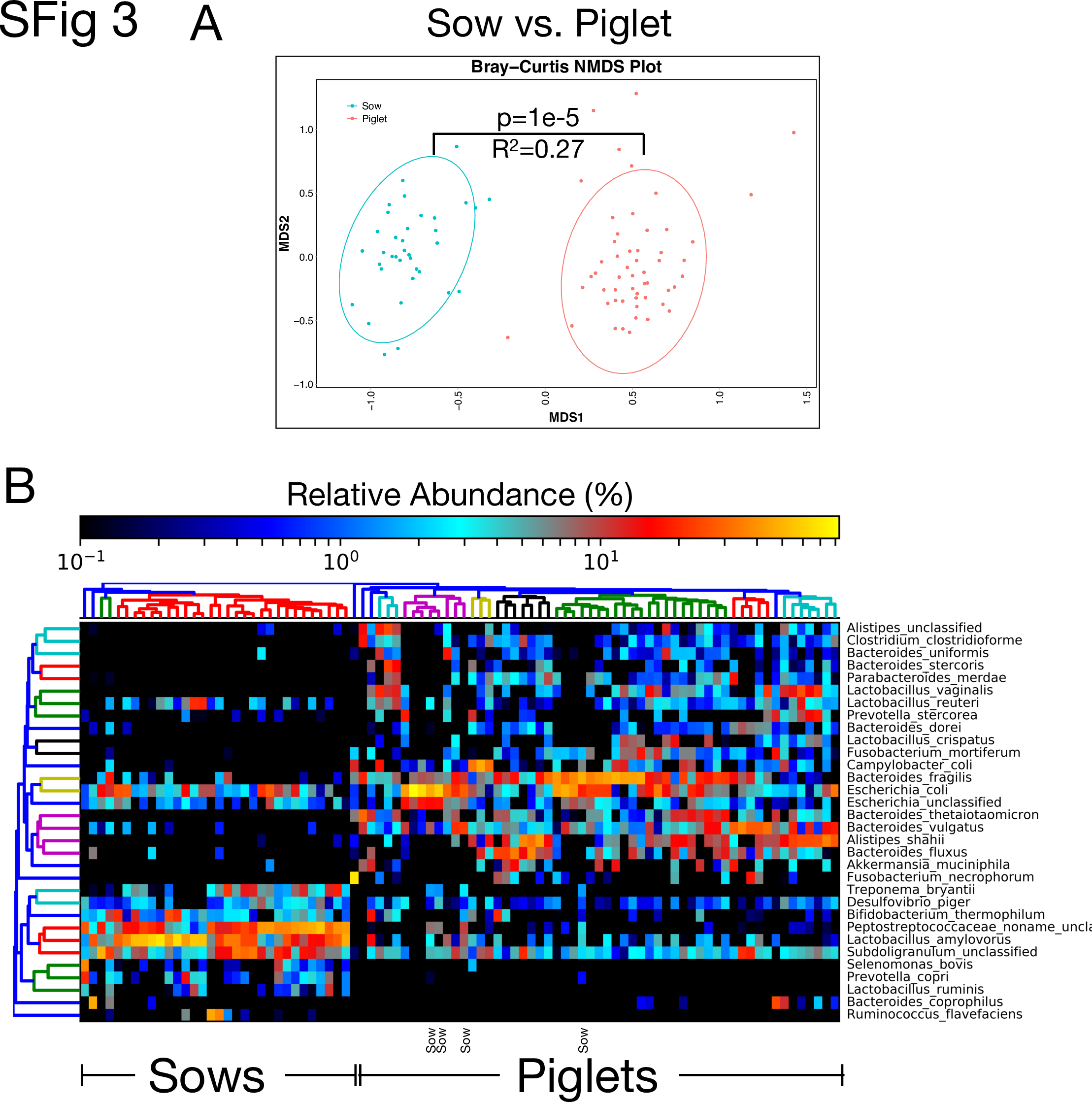

### Supplemental figure 4

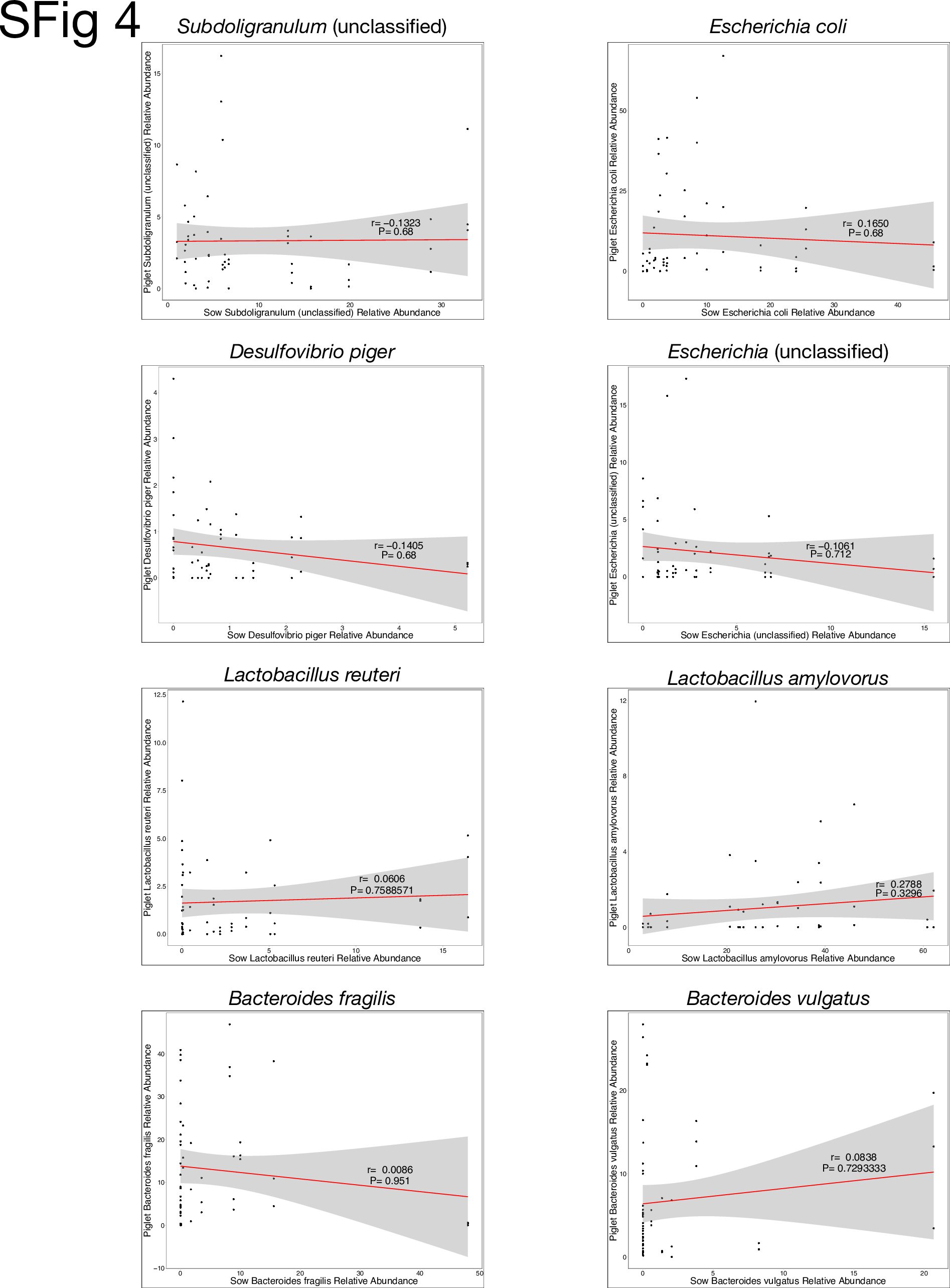

### Supplemental figure 5

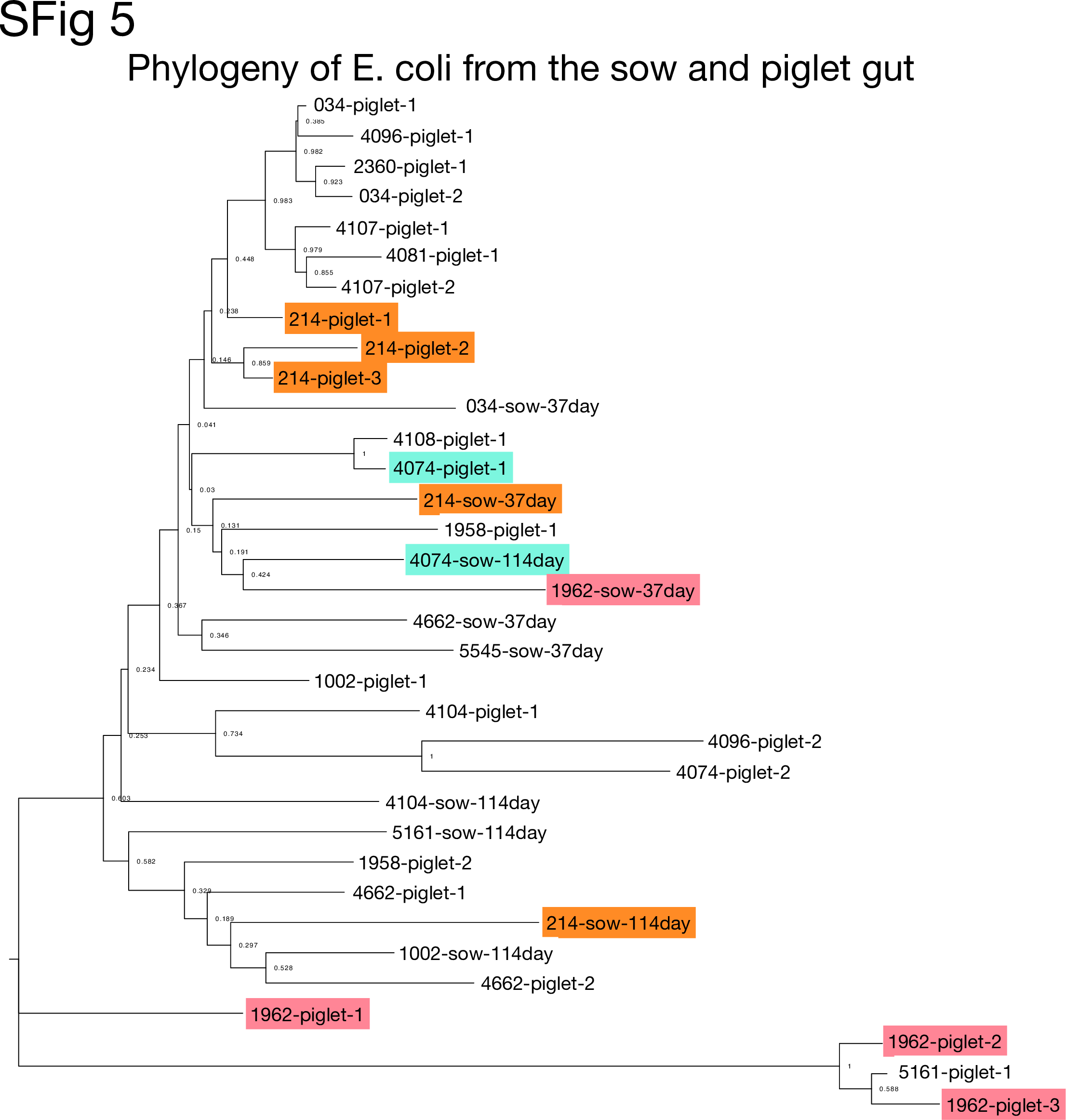
